# What is a cognitive map? Organising knowledge for flexible behaviour

**DOI:** 10.1101/365593

**Authors:** Timothy E.J. Behrens, Timothy H. Muller, James C.R. Whittington, Shirley Mark, Alon B. Baram, Kimberley L. Stachenfeld, Zeb Kurth-Nelson

## Abstract

It is proposed that a cognitive map encoding the relationships between entities in the world supports flexible behaviour, but the majority of the neural evidence for such a system comes from studies of spatial navigation. Recent work describing neuronal parallels between spatial and non-spatial behaviours has rekindled the notion of a systematic organisation of knowledge across multiple domains. We review experimental evidence and theoretical frameworks that point to principles unifying these apparently disparate functions. These principles describe how to learn and use abstract, generalisable knowledge and suggest map-like representations observed in a spatial context may be an instance of general coding mechanisms capable of organising knowledge of all kinds. We highlight how artificial agents endowed with such principles exhibit flexible behaviour and learn map-like representations observed in the brain. Finally, we speculate on how these principles may offer insight into the extreme generalisations, abstractions and inferences that characterise human cognition.

## Introduction

In the last two decades and more, computational and behavioural neuroscientists have found formal explanations of neural signals that control behaviour in carefully controlled repetitive scenarios (e.g. (Behrens et al., 2007; Daw et al., 2006; O’Doherty et al., 2004; Platt and Glimcher, 1999; Schultz et al., 1997)). In some instances, these models predict neuronal activity with truly exquisite precision (Cohen et al., 2012; Gold and Shadlen, 2007; Schultz et al., 1997) and when paired with heavy computational resources, related algorithms have had extraordinary successes in training artificial agents to super-human levels in games as diverse as Atari (Mnih et al., 2015) and Go (Silver et al., 2016). However, there is a stark gap between the types of behaviour these models can account for and the sophisticated inferences that characterize much of human behaviour. Human and animal behaviour is flexible. We can choose how to act by exploiting actions that have worked in the past, but also based on experiences that are only loosely related; we can imagine the consequences of entirely novel choices. We can abstract important features of experiences and generalise them to new situations. These differences were clearly articulated by Tolman as he watched rats make flexible inferences in complex mazes. They would learn rich details of the mazes in the absence of any rewards and to the benefit of future behaviour. For example, after unrewarded exposure to mazes, rats would take short-cuts to reach rewards (Tolman and Honzik, 1930) or would find new routes when old ones were blocked (Tolman et al., 1946). Such behaviours inspired Tolman to coin the term ‘cognitive map’ referring to a rich internal model of the world that accounts for the relationships between events, and predicts the consequences of actions.

For Tolman, this cognitive map was a systematic organisation of knowledge that spanned all domains of behaviour (Tolman, 1948). However, its biggest influence in cognitive neuroscience has been in the study of spatial behaviours (O’Keefe and Nadel, 1978), perhaps because the literal interpretation of the term ‘map’ gives clear predictions of neural activity. Even Tolman cannot have imagined the beautiful precision with which map-like representations are reflected in the activity of single neurons in the hippocampal-entorhinal system (Figure 1). The most celebrated of these neurons are active at particular places in the map. ‘Place’ cells in the hippocampus restrict their activity (usually) to a single location in space (O’Keefe and Nadel, 1978). ‘Grid’ cells in the medial entorhinal cortex fire at multiple place fields equally placed on a triangular grid (Hafting et al., 2005), and are therefore able to represent vector relationships and distances between different spatial locations (Bush et al., 2015; Stemmler et al., 2015). Along with these come a veritable zoo of less celebrated but equally remarkable cells that reveal how ‘knowledge’ is organised in the map (Grieves and Jeffery, 2017), such as band cells (Krupic et al., 2012), and cells that encode the vector relationships to borders (Solstad et al., 2008), objects (Høydal et al., 2018), rewards (Gauthier and Tank, 2018) and goals (Sarel et al., 2017), alongside cells that encode the current head direction (Taube et al., 1990); or cells that encode the locations of other agents on the map (Danjo et al., 2018; Omer et al., 2018).

**Figure 1:**
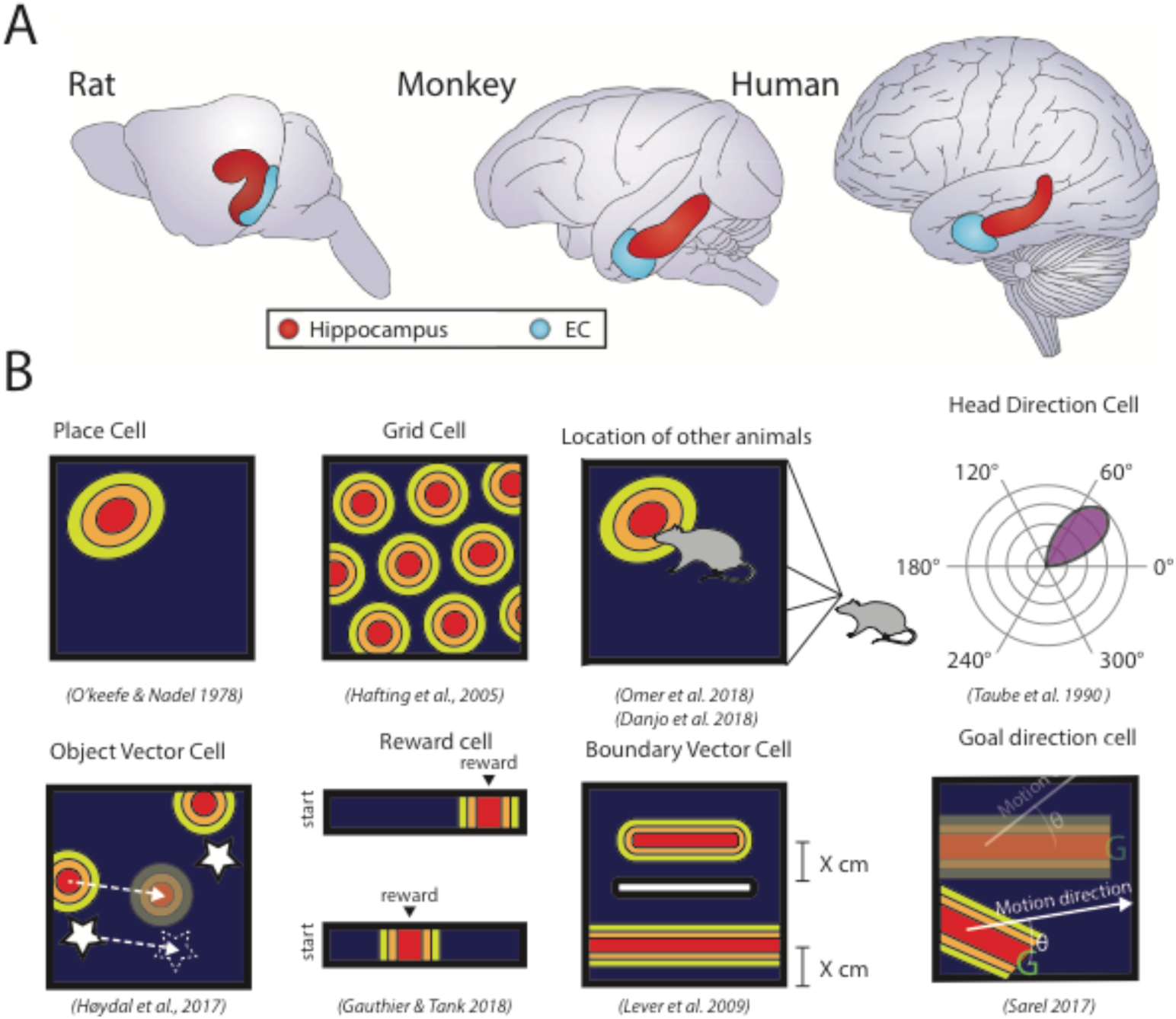
The hippocampal zoo. A) Anatomical location of the hippocampus and entorhinal cortex in different species. Adapted from (Strange et al., 2014). B) A variety of cells in the hippocampal formation represent different spatial variables. Place cells (O’Keefe and Nadel, 1978) are active when an animal is in a single (sometimes multiple) location. Grid cells (Hafting et al., 2005) are active when an animal is in one of multiple locations on a triangular lattice. “Social place cells” (Danjo et al., 2018; Omer et al., 2018) are active in one animal when it observes another animal is in a particular location. Head-direction cells (Taube et al., 1990) are active when an animal’s head is facing a particular direction. Object-vector cells (Høydal et al., 2018) are active when an animal is in a particular direction and distance from any object. Reward cells (Gauthier and Tank, 2018) are active when an animal is in the vicinity of reward. Boundary vector cells (Lever et al., 2009) are active at a given distance away from a boundary in a particular allocentric orientation. Goal direction cells (Sarel et al., 2017) are active when the goal of an animal is in a particular direction relative to its current movement direction. The green ‘G’ indicates the goal location.

These spatial cells appear to have specialised functional representations such that each plays an important role in understanding and navigating a 2D world. Notably, however, the same brain structures containing these cells play important roles in neural processes that relate to a broader view of a cognitive map, such as generalisation, inference, imagination, social cognition, and memory (Hassabis et al., 2007; Van Der Meer et al., 2012; Ólafsdóttir et al., 2015; Tavares et al., 2015). It is therefore a challenge to understand how such cells might help us organise knowledge in the complex high-dimensional non-spatial cognitive map that Tolman envisaged. In this review, we show how recent work is beginning to find unifying explanations for these apparently disparate functions by looking to ideas from reinforcement learning and statistical learning, and investigate whether such formalisms may not only explain neuronal responses in spatial tasks, but also provide opportunities for the types of powerful inferences and generalisations of structural knowledge that underlie flexible human behaviour.

### Organising structural knowledge for flexible learning

Whilst Tolman was watching rats running in mazes, another hero of psychology, Harlow, was asking human and non-human primates to choose between two stimuli to find a reward. Discriminating a rewarding from an unrewarding stimulus should not require any sophistication or flexibility at all – the animal could just preferentially repeat rewarded choices – but Harlow noticed something interesting. As subjects had more and more experience of the task (with different stimuli each time) they got better and better at learning new discriminations (Figure 2A-B). As well as learning which was the better stimulus, the subjects were learning something abstract about how to perform the discrimination. He termed this abstract knowledge a ‘learning set’ (Harlow, 1949). In this and the following sections, we will contend that acquiring such a learning set requires an abstract representation of the structure of the task that encodes relationships between task events. This kind of representation allows inferences from remote observations, and generalisation of information across different tasks with similar structure. We will also argue that grid cell activity is an example of such a representation in the spatial domain, such that grid cells encode statistical regularities in spatial navigation that occur due to the common structure of all two-dimensional spaces.

**Figure 2:**
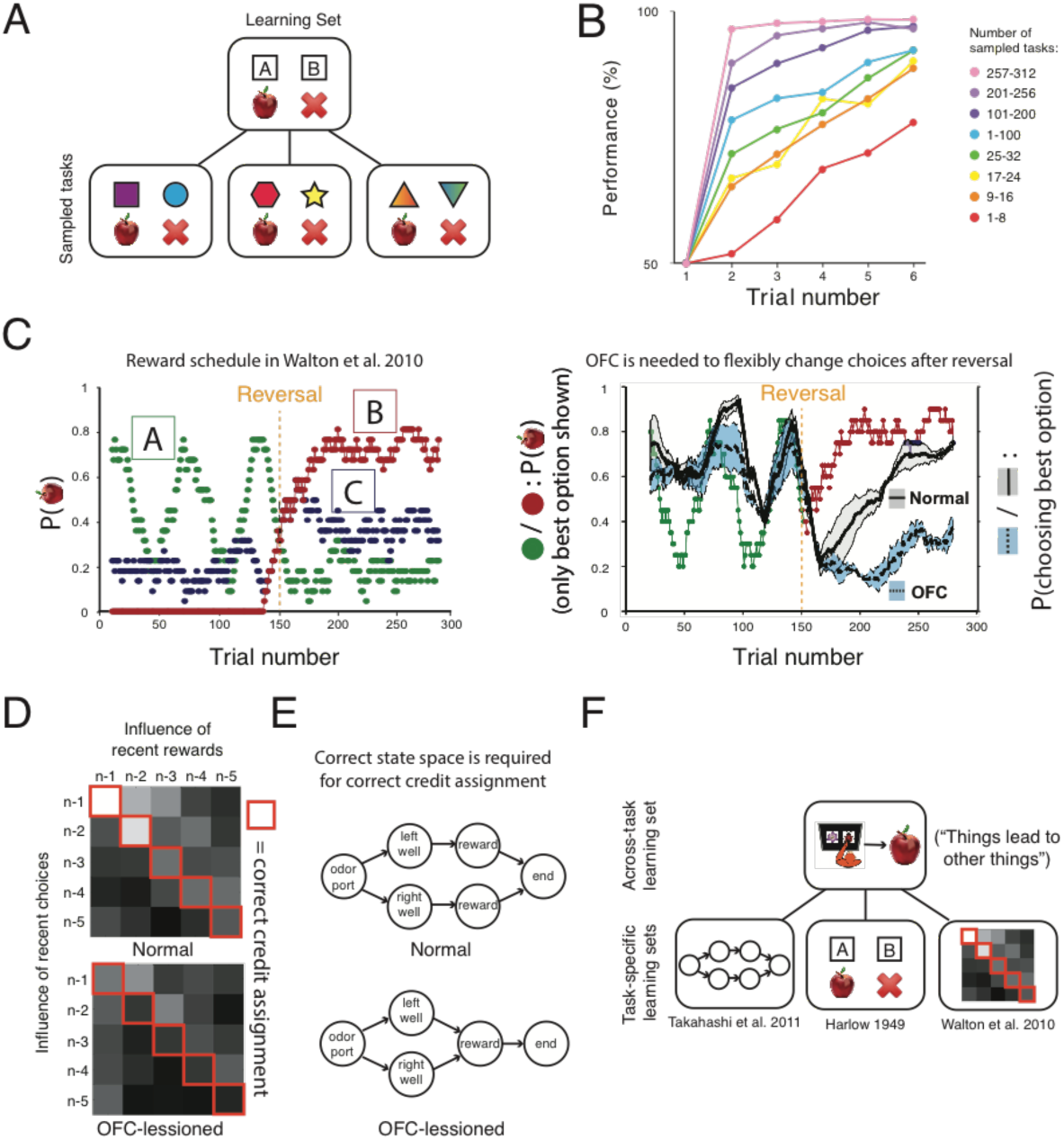
The importance of learning sets - Harlow and beyond. We will describe three tasks relying on learning sets. (Harlow, 1949) demonstrates the learning of task structure from repeated exposure to the same task. (Walton et al., 2010) and (Takahashi et al., 2011) demonstrate the implications of OFC-lesions on a part of the learning set common across many different tasks. A) Schematic of Harlow’s task. Different instantiations of the task share a common underlying structure (“only one object is rewarded”) that can be exploited to facilitate faster learning. B) Accuracy data from (Harlow, 1949). Over multiple exposures to the task, monkeys acquire this underlying structure, termed a “learning set”, and use it to learn faster in new instantiations of the task. C) Monkeys can also learn to track changing probabilities of reward, but OFC lesions cause an intriguing deficit whereby they can track changes in reward unless the best option reverses. D) This is because they no longer know which recent choice caused which recent reward. In these regression plots, correct attribution between choice and reward is on the diagonal. OFC lesioned animals learn using the off-diagonal terms which imply that if average recent reward is high (low) the animal increases (decreases) their preference for their average recent choice, ignoring which choice caused which reward. C) and D) Adapted from (Walton et al., 2010). E) Takahashi et al. showed a similar effect in rodents. (Wilson et al., 2014) demonstrated that it could be accounted for by a model in which the animal could not distinguish latent states (reduced state diagram shown). Here the states “reward after left choice” and “reward after right choice” are represented separately in the model that best accounts for control data. The fact that they are not in the model that best accounts for OFC-lesioned data reflects the rodents’ inability to pair the reward with the choice that caused it. We refer the reader to their beautiful paper for other examples of state space-deficits caused by OFC lesions. F) A learning set for a task is composed not only from the abstract structure common to different realisations of a specific task (as demonstrated in Harlow’s findings), but also from basic knowledge which is generalisable to many tasks. For example, the notion of contingency – that a choice leads to the immediately following outcome - is a feature of many different tasks and life experiences. This part of the learning set underlies the results in (Walton et al., 2010) and (Takahashi et al., 2011). The full learning set model might consist of many higher-order across-task learning sets, converging to different degrees on each task-specific learning set.

To acquire such a learning set, you need to learn abstract relationships between different stimuli, such as “if one stimulus is rewarded, the other is not”, or “the rewarded stimulus might change after a number of trials”. Part of this learning process includes learning basic knowledge about how the task works. For example, you need to know the reason you are getting a reward now is because of the stimulus you just chose, and not the stimulus from 3 trials ago or the door that just opened in the corridor. When lesions are made to the ventral prefrontal cortex (vPFC) (orbitofrontal and ventrolateral prefrontal cortex^1^) in macaque monkeys, this ability is abolished (Rudebeck and Murray, 2011; Walton et al., 2010). Animals no longer assign the credit for each reward to the contingent choice that caused it, but instead to an imprecise running average of recent choices. After vPFC lesions, macaques that once knew the structure of the task, now learn by smooth temporal correlations (Walton et al., 2010) (Figure 2C-D). Whilst such a strategy can work well in stable environments, it leads to disastrous performance when behavioural flexibility is required, such as when reward contingencies change. These vPFC properties are not unique to the brains of macaque monkeys. In humans, fMRI signals in vPFC reflect precise task contingencies when other reward signals in the brain do not (Jocham et al., 2016). In rodents, if unilateral lesions are made to OFC, dopaminergic cells in the same hemisphere continue to report a veridical reward prediction error, but with a prediction that no longer reflects which choice has caused the reward (Takahashi et al., 2011).

What does it mean to have a representation of the structure of a problem? Thinking about these issues more formally has led to a richer understanding (Wilson et al., 2014). In reinforcement learning, such behavioural control problems can be cast in terms of trying to find a *policy* that will maximise long term cumulative reward. The problem is characterised by states, *s*, and probabilistic transitions between states, *P(s’*|*s*, *a*), which may be controlled by actions, *a*. The policy, *π*, determines the probability of choosing each action in each state; *π = P*(*a*|*s*). If *r* denotes the instantaneous reward received at a current state, *V_π_* denotes the expected cumulative reward over the foreseeable future under a policy, and *γ* the discount factor weighting immediate rewards higher, then, after some maths, our goal becomes finding a policy that maximises value (the following equation):

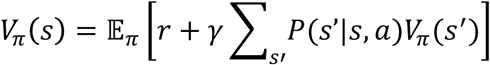

In this framework, the burden of representing the problem structure is carried by the state definition, *s*, and the transitions, *P(s’*|*s, a*) (Box 1). These respectively describe how is the task divided up into different elements – for example, the state of having just seen a particular stimulus – and how one element leads to another. Wilson and colleagues showed that the credit assignment deficits described above, along with other types of deficit commonly observed after ventral prefrontal lesions, are predicted by a reinforcement learning agent that only learns from immediate sensory observations, and does not assign credit to abstract states (such as “I just chose stimulus A”, Figure 2E). It is argued, then, that activity in the OFC must encode the current location in a latent, or unobserved, state space. Indeed, this exact information can be decoded from the OFC fMRI signal when humans engage in a complex task with a well-defined state space (Schuck et al., 2016).

To account for effects of learning set, however, a state representation must do more than simply label the current state. First, it must encode how this state relates to other states in the world (so an animal can know, for example, that if one state is not rewarded the other state likely is, or if their spouse’s wallet is on the table then they are more likely in the garden than the pub). Second, it must encode states in a fashion that generalises across different sensory realisations of the task. A key feature of both Harlow’s experiments and the OFC tasks in (Walton et al., 2010) is that every example of the task used different stimuli, but animals improved on the task nevertheless. Indeed, when monkeys are asked to make economic choices between different amounts of two juices A and B, OFC neurons encode a rich variety of value and task-related variables (Padoa-Schioppa and Assad, 2006), but when these two juices are replaced with two different juices C and D, the same neurons encode the exact same variables for the new juices (Xie and Padoa-Schioppa, 2016) (Figure 3C-D). Indeed, lesions to neighbouring vlPFC cause particular deficits in generalising knowledge between different stimulus sets (Rygula et al., 2010).

**Figure 3:**
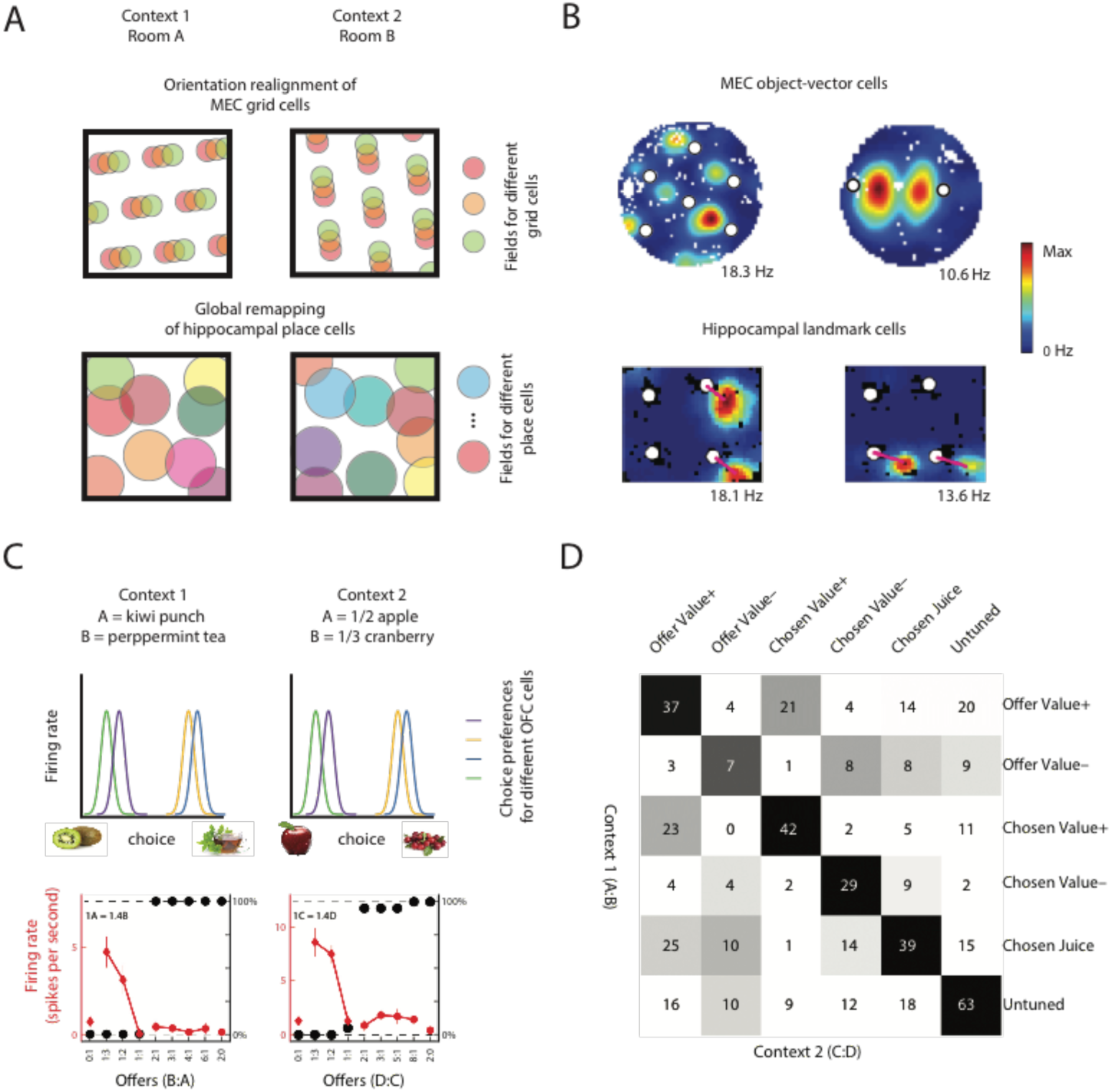
MEC and OFC neurons generalise across different contexts. A) In a spatial remapping experiment, animals are moved between two different environments. Entorhinal grid cells maintain a constant spatial phase structure (top), in contrast with the global remapping of hippocampal place cells (bottom) (Bostock et al., 1991; Leutgeb et al., 2005). B) MEC object vector cells respond specifically when an animal is at a given direction and distance from any object, regardless of identity of the object (top). Cells with object-vector properties are also found in the hippocampus (bottom). These cells, however, respond to only a subset of the objects (bottom). MEC data from (Høydal et al., 2018), Hippocampal data from (Deshmukh and Knierim, 2013). C) When monkeys choose between pairs of different juices offered in different amounts, OFC neurons encode the same decision variables across contexts, such that two neurons ‘supporting’ the same decision for one pair of juices also support the same decision in different pairs of juices (top). Firing rate of an example OFC neuron as a function of the quantity ratio of the two juices (bottom). Black symbols represent the percentage of trials in which the monkey chose the less preferred juice. Red symbols represent the average firing rate. D) Population analysis demonstrating OFC neurons generalise over different contexts. Each task-related OFC neuron was classified according to its strongest encoded decision variable (offer value, chosen value, chosen juice) and the sign of this encoding. This was done in two separate contexts (A:B choice / C:D choice shown in rows/columns). Numbers are cell counts for encoding a pair of variables in the two contexts. Statistical significance was assessed via a null distribution (greyscale colors are P-values). The strong diagonal indicates neurons tend to encode the same variable with the same sign in both contexts. (C) and (D) adapted from (Xie and Padoa-Schioppa, 2016).

### Generalising spatial representations

So far, we have argued that vPFC representations in learning tasks have three important computational properties. (1) They encode ‘location’ in a state space of the task. Their representation of location (2) embeds structural knowledge of the relationships between different states, and (3) can be generalised across tasks with shared statistical structure but different sensory events. These three features of OFC activity resemble the key computational properties of entorhinal cells in spatial domains. Grid cells famously encode location in a spatial task. They do so with a representation that has implicit within it knowledge of the spatial relationships between all locations, allowing remote inferences (Bush et al., 2015; Stemmler et al., 2015) (for further convincing see next section), and this structural code generalises across different sensory environments. To understand this final point, consider what happens in a hippocampal remapping experiment, in which animals are moved between two different boxes (for example, with different wall colours). In these experiments hippocampal place cells remap when the sensory environment changes (Bostock et al., 1991; Leutgeb et al., 2005). Neighbouring place cells in one environment are unlikely to be neighbours in the other. By contrast, except for a rigid body change, grid cells do not remap between environments. Within a module, phase neighbours in one environment are phase neighbours in the other (Fyhn et al., 2007) (Figure 3A). The entorhinal representation of location therefore embeds the structural information about general relationships in 2D space that is common amongst all environments. Similarly, object vector cells in entorhinal cortex activate for any object present in the environment (Høydal et al., 2018), whereas similar cells in hippocampus activate for only a subset of objects (Deshmukh and Knierim, 2013) (Figure 3B).

With these relationships in mind, it is intriguing that deficits in learning set can be achieved by transection of the fornix (which disconnects frontal cortex from the hippocampus) (M’Harzi et al., 1987) or by fronto-temporal disconnection (Browning et al., 2006). Similarly, interactions between OFC and hippocampus appear to be important for correctly updating task representations in humans (Boorman et al., 2016) and rodents (Wikenheiser et al., 2017). It is therefore interesting to understand how we can relate these more general forms of behaviour to hippocampal representations familiar from spatial navigation tasks.

These relationships may be more than simply theoretical. By casting reinforcement learning problems in continuous but non-spatial domains, recent studies suggest that place and grid cells may have a broader role than the coding of physical space. These studies show activity of cells in the same regions as place and grid cells, measured either with fMRI or directly, code for non-spatial information in a manner analogous to the coding of spatial information.

In humans, it is possible to record grid-like activity during virtual reality navigation either directly from the activity of entorhinal cells during surgery (Jacobs et al., 2013) or indirectly with fMRI (Doeller et al., 2010). Whilst fMRI cannot give access to cellular activity directly, the hexagonal symmetry of the grid pattern has a striking shadow in the fMRI signal. As subjects move in the VR environment, fMRI activity exhibits a 6-fold oscillation as a function of running direction (Doeller et al., 2010) (Figure 4A-B). Notably, this pattern can be observed not only when subjects navigate in virtual spatial worlds, but also when they are engaged in an operant non-spatial task that has the same statistical structure as space (2D continuous organisation) (Constantinescu et al., 2016). Instead of moving in space, subjects watch as a cartoon bird morphs in two dimensions (the lengths of the neck and legs). Their task is to predict when these birds will match the appearance of one of several target birds that are associated with different rewards. The instantaneous change in the appearance of the bird describes a vector in a 2D conceptual space defined by the neck- and leg-lengths, and grid-like coding is inferred by looking for a 6-fold oscillation in fMRI activity as a function of this vector. This pattern can be observed in entorhinal cortex but also in other brain regions including ventral frontal cortex (Constantinescu et al., 2016) (Figure 4C). By contrast, in hippocampus, cells fire to specific abstract stimuli, such as Jennifer Aniston (Quiroga et al., 2005), and therefore code information in a fashion analogous to place cells in spatial domains.

**Figure 4:**
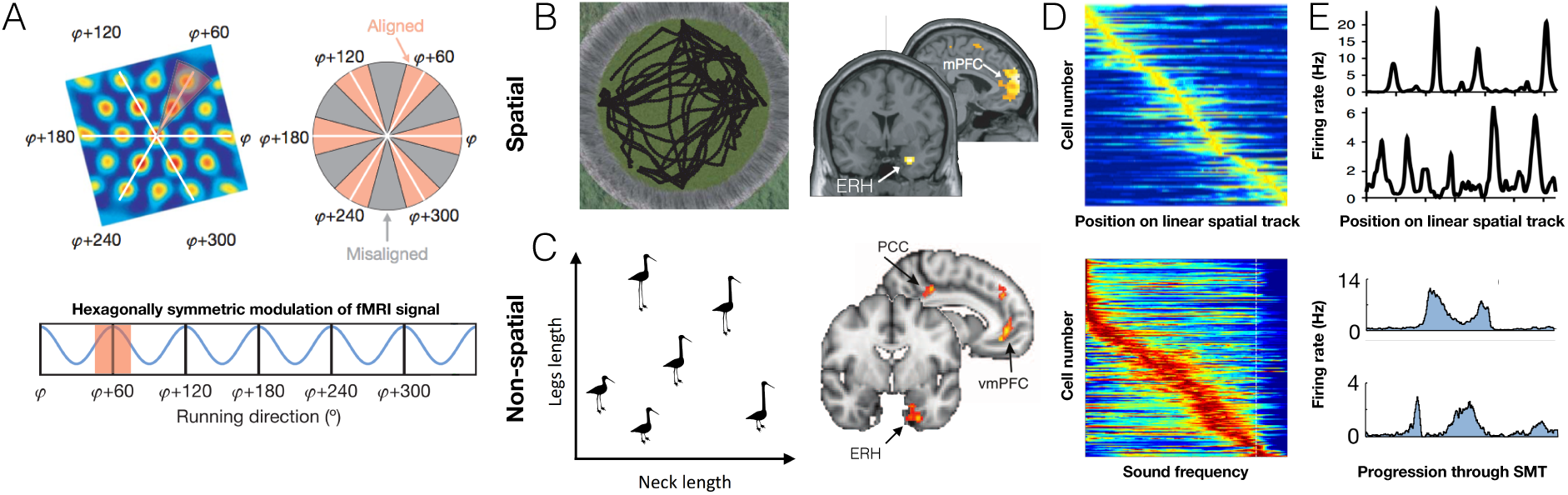
Generalising spatial representations. A) Logic behind measuring grid cells with fMRI. Trajectories through a 2-dimensional space can be either aligned or misaligned with the axes of the grid code (white lines denote grid axes). Greater signal for trajectories aligned versus misaligned with the grid results in a hexagonally symmetric sinusoidal modulation of the fMRI signal with movement direction. Adapted from (Doeller et al., 2010). B) This modulation of the fMRI signal – providing evidence for grid cells – is observed in entorhinal cortex (ERH) when participants navigate through virtual reality spatial worlds (Doeller et al., 2010). Left is an aerial view of the spatial task map. C) The same signal is observed when participants navigate through an abstract conceptual space defined by two continuous dimensions: the neck length and leg length of “stretchy birds”, suggesting grid cells code for non-spatial dimensions (Constantinescu et al., 2016). D) Cells in the hippocampus fire at specific sound frequencies in a non-spatial, sound manipulation task (bottom) in a manner analogous to the representation of spatial locations in place cells on a linear track (top). In both panels, each horizontal line shows normalised activity of one cell, as a function of either distance along a linear spatial track or sound frequency. Cells are ordered according to the position of the firing field. Top adapted from (Miao et al., 2015) and bottom from (Aronov et al., 2017). E) Grid-like coding in entorhinal cortex of a linear spatial track (top) and progression through the sound manipulation task (bottom). Top adapted from (Yoon et al., 2016) and bottom from (Aronov et al., 2017).

Similarly, for rodents, the task of “holding a lever whilst a tone increases in frequency, and then releasing it at a target frequency to get a reward” is not obviously a spatial one, but it has a topology familiar from space - one frequency leads to the next in the operant box in the same way as one place leads to the next on a linear track. When rodents perform this task (Aronov et al., 2017), hippocampal cells exhibit place-like firing fields, but for different frequencies rather than places, and entorhinal cells (including spatial grid cells) exhibit multiple distinct fields at different frequencies that resemble the firing fields of grid cells on a linear track (Yoon et al., 2016) (Figure 4D-E). Grid cells also encode gaze location on a 2D image in both nonhuman (Killian and Buffalo, 2018) and human primates (Julian et al., 2018; Nau et al., 2018).

There is evidence, then, that place and grid patterns are neither unique to spatial navigation nor, in humans at least, unique to the hippocampal formation (see also (Jacobs et al., 2013)). Instead they may reflect the 2D topology that is inherent to space, as well as characterize other domains. These results suggest the role of place and grid cells in spatial cognition is a specific instance of more general coding mechanisms realised in the hippocampus and connected regions.

### Unifying spatial and non-spatial coding under a common framework

In order to understand more formally what this means, it is useful to return to the RL framework laid out in the previous section (and in Box 1). Because RL is a general framework, it is not limited to explaining operant tasks. It can equally easily be used to give a fresh perspective on the spatial navigation problem (Gustafson and Daw, 2011). For example, consider a rat running on a linear track. Using the RL framework, we can express this task by choosing our states *(s)* to be the different locations along the track (Figure 5A). The probability that the rat moves from state to state depends on its policy and if given by 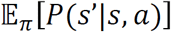, however if we ignore the effects of policy and assume that all movement is through diffusion (random walk on the graph), then this same equation tells us the environment’s transition probabilities between states (**T**). This matrix effectively tells us which states are neighbours and therefore on a linear track encapsulates the 1-dimensional topology of the problem space (Figure 5A). It is a useful matrix to know. If you are planning your future and want to know where you will likely be at the next time-step, you can simply multiply your current state vector, **s**, by **T** to give **Ts**. In two time-steps, your state probability distribution will be **T^2^*s***, and in 3-steps **T^3^*s*** etc (Figure 5A).

**Figure 5:**
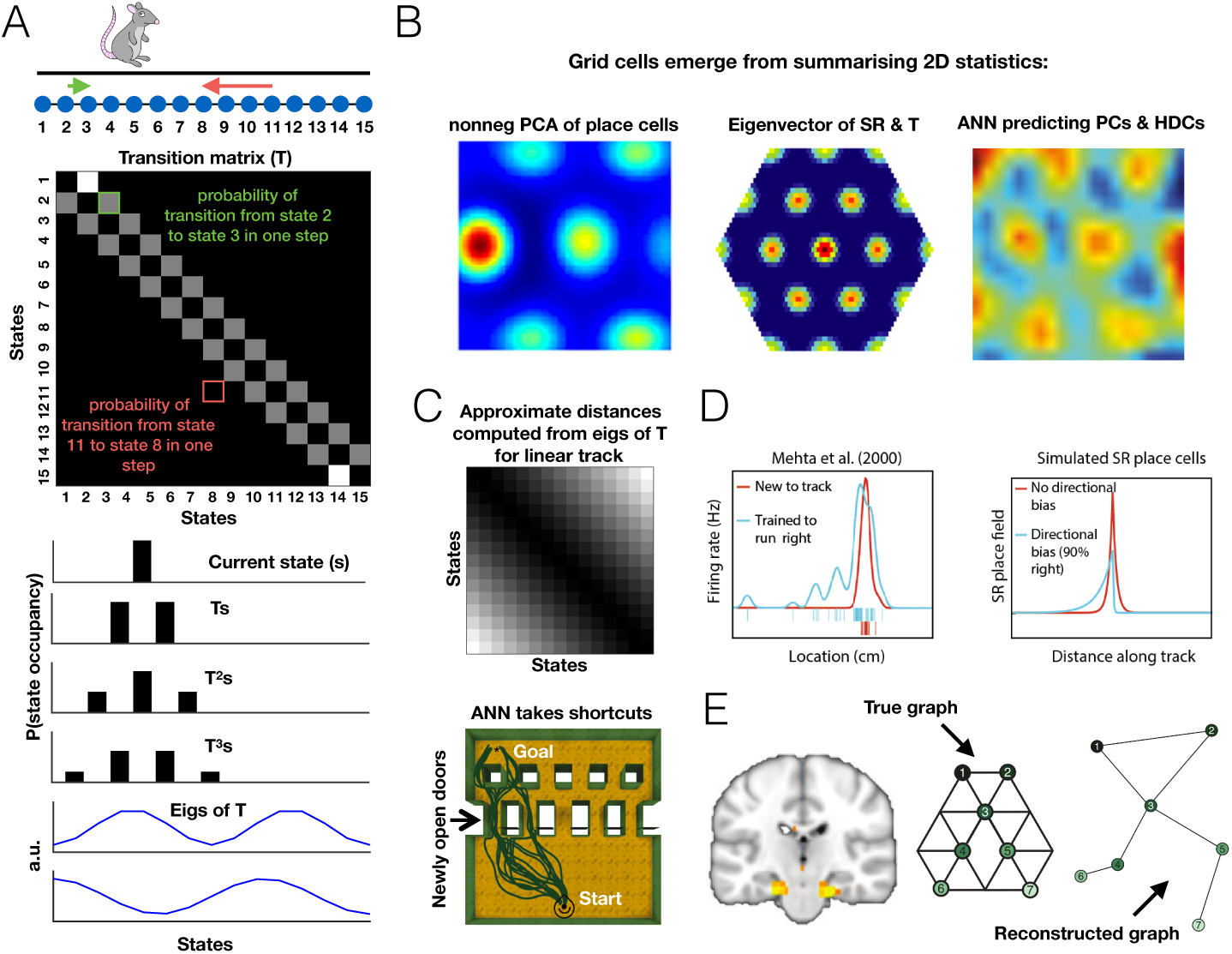
Unifying spatial and non-spatial representations under a common framework. A) Reinforcement learning can offer a fresh perspective on spatial cognition by considering locations as states. For example, a linear track can be thought of as a series of neighbouring states. This 1D topology can be represented in a states by states transition matrix **T**, with element T*_i,j_* corresponding to the probability of transitioning from state *i* to state *j*. Thus, this matrix represents structure. Indeed, multiplication of the current state, **s**, by **T^n^** gives a distribution over state occupancy n steps into the future. Notably, eigenvectors of this transition matrix are periodic, suggesting they contain non-local knowledge about the structure that may be useful for computing distances (see D). B) Grid cell-like firing fields can be obtained by casting 2D space under this state space framework. The eigenvectors of the covariance of 2D-distributed place cells (obtained by nonnegative principal component analysis, PCA), of 2D transition matrices, and of successor representations of 2D state spaces are all grid-like, as are units of an artificial neural network (ANN) tasked with predicting 2D-distributed place cells (PCs) and head direction cells (HDCs). Figures adapted from (Dordek et al., 2016) (left), (Stachenfeld et al., 2017) (middle) and (Banino et al., 2018) (right). C) These grid-like representations embed structural knowledge of the relationships between states. Since the eigenvectors of **T** are also the eigenvectors of the successor representation, they can be used to compute approximate distances between all pairs of states, which can be used to facilitate planning (top: element *I,j* of the matrix is the distance between state *i* and *j;* lighter colour denotes larger distance). Grid codes provide a basis for vector-based navigation (Bush et al., 2015), allowing the ANN with grid-like units to take shortcuts (bottom) (Adapted from (Banino et al., 2018)). D) Using representations that, rather than encoding current location, are predictive of successive states (successor representation), it is possible to explain policy dependent phenomena in the hippocampal formation. For example, if an animal moves only one way down a linear track, successor representations skew towards the start of the track to predict their future state, as observed in hippocampal place cells. Figure adapted from (Stachenfeld et al., 2017). E) This same framework can explain the neural representation in hippocampus and entorhinal cortex of a discrete, non-spatial state space. The true underlying graph structure of this state space can be reconstructed from the neural activity, suggesting these regions represent discrete as well as continuous tasks. Adapted from (Garvert et al., 2017).

This type of future-thinking is an example of model-based reinforcement learning, where an agent that knows the states and transitions (and therefore has a ‘model’ of the world) can simulate its future step-by-step and make decisions about which future is best (Daw et al., 2005, 2011; Sutton and Barto, 1998). Whilst there is strong evidence for this type of future projection at choice points in spatial tasks (Doll et al., 2015; Johnson and Redish, 2007), it is very different in spirit to how we think about most navigation problems. In navigation problems, instead of sequentially planning through each neighbouring location, agents are able to use the known Euclidian properties of 2D space to infer local vectors that will connect distant points. Can similar inferences be made in RL state-spaces? Without delving deep into the mathematics, there *is* a set of vectors from which it is possible to compute all of the n-step transition matrices (**T**,**T**^2^,**T**^3^ **… T^n^…**) by simple linear combination. These are the eigenvectors of **T** (Figure 5A). These vectors linearly encode all futures and from them it is easy to compute distances between any pairs of states without the need for expensive step-by-step simulation (Baram et al., 2017; Stachenfeld et al., 2017). For continuous worlds these eigenvectors are periodic and for 2D worlds, they have grid-like properties (Dordek et al., 2016; Stachenfeld et al., 2017) (Figure 5B).

It is notable that, because place cells index overlapping states in a 2D world, these eigenvectors are also the principal components of place cell activity (Dordek et al., 2016; Stachenfeld et al., 2017). The eigen/grid code can therefore also be thought of as a code that captures the variance in the place-cell population in an information-efficient manner. Indeed, when a recurrent neural network is trained to predict place cell (and head direction cell) activity (Banino et al., 2018), or location (Cueva and Wei, 2018), as it navigates around an open field, it develops grid-like cells as its preferred linear representation (Figure 5B). By using these cells in more complex environments, it can solve navigation problems that require vector navigation (such as finding short-cuts that have never previously been taken) (Banino et al., 2018) (Figure 5C).

To date, we have been considering how to represent likely future experiences in situations where the world has useful structure but behaviour is random. In fact, because transitions depend on choices, the expected transition probabilities, **T**, change if the animal shows statistical regularity in its behaviour (for example, if the animal likes to approach food-sources). This *policy-dependence* can be harnessed to make predictions about hippocampal representations that are not immediately obvious from spatial considerations alone. To do this, it is necessary to re-envisage the place cell representation. Instead of coding where the animal is now, once an animal becomes familiar with an environment, it is possible to encode its best estimate of where it will be in the imminent future. This is clearly a useful representation for controlling behaviour, as it allows the animal to rapidly evaluate which local choices have profitable futures. In reinforcement learning, this is termed a *successor representation* (Dayan, 1993) as it predicts the expected *successor states*. By assuming that the place cells encode these successors rather than current location, it is possible to account for a number of seemingly disparate findings in the place cell literature, such as the tendency of place fields to stretch slowly towards the start of a 1-way linear track or to clump in high densities around rewarding locations (Stachenfeld et al., 2017) (Figure 5D).

This general framework goes some way to explaining why place and grid representations are not unique to spatial situations, but also implies that they should not be unique to continuous environments. Whilst this is an ongoing topic of investigation, there is suggestive evidence that it will be a profitable one. In humans, FMRI similarity measures in hippocampus (Garvert et al., 2017; Schapiro et al., 2013; Stachenfeld et al., 2017) and entorhinal cortex (Garvert et al., 2017) respect the statistical transitions of discrete state-spaces even when subjects are unaware that transitions are non-random. By examining these representations it is possible to reconstruct directly from the neural data state distance matrices that resemble the true transition or successor distances between states (Figure 5E). In operant tasks, rather than simulating all possible transitions online, there is evidence human behaviour relies on precompiled transition distances consistent with the successor (or eigen-) representation (Momennejad et al., 2017a) and that these precompiled distances rely on offline activity in hippocampus and ventral frontal cortex (Momennejad et al., 2017b). Similarly, in a rodent operant task, a manipulation of either hippocampus or orbitofrontal cortex prevents the animals from using the state transition structure to guide their next choice (Miller et al., 2017, 2018).

### Inferences, abstractions and factorised representations of tasks

Within either a reinforcement learning task or a spatial environment, then, a clever representation of how states are related can allow for flexible inferences. But how does the brain come by such a representation each time it encounters a new problem? In this section, we argue that structural knowledge can be abstracted away from its sensory inputs and therefore generalises to new environments, state spaces and tasks. We argue that some structural representations are broadly required across many tasks that require flexible inferences and rapid learning.

As discussed, such inferences are routinely made by animals in the spatial domain. One way to understand how an animal might take a new shortcut, for example, is to consider that the statistical structure of 2D space places strong constraints on what state transitions are possible. When the animal is moving in a spatial environment, it samples some of those states and transitions, and can use this *prior structural knowledge* to fill in many states and transitions that it has not seen, but are implied by the 2D nature of the problem. Are there other situations in nature where similar constraints might apply? Are there non-spatial scenarios where humans and animals can make structural inferences with no prior experience of a problem?

To find them, we don’t have to look far, either in topological complexity or in neural anatomy. If animals are independently taught that they should choose stimulus A over B and B over C, they will infer that they should choose A over C on first presentation – a phenomenon known as transitive inference (Burt, 1911; Dusek and Eichenbaum, 1997; von Fersen et al., 1991; Mcgonigle and Chalmers, 1977) (Figure 6A). In primates, such lists can be long (e.g. ABCDEFG), and can be flexibly reconfigured – for example stitched together by presentation of the critical link (Treichler and Van Tilburg, 1996). Similarly, animals can stitch together temporally distinct episodes into a linear time representation, and use this representation to make causal inferences. In a sensory preconditioning paradigm, for example, animals are taught that A leads to B and later that B leads to reward (Jones et al., 2012). When they are later asked to choose between A and a control, they choose A – the stimulus that implies a path to reward. Whilst it is possible to solve both of these tasks by simpler associative mechanisms, in the case of transitive inference, at least, evidence strongly suggests that animals instead rely on abstract knowledge of the linear structure (Gazes et al., 2012; Jensen et al., 2015, 2017; Lazareva and Wasserman, 2012). For sensory preconditioning, the jury is still out in animals, but any reader who was able to decipher the plots of “Pulp Fiction” or “Kill Bill” from the sporadic, interleaved, and often time-reversed episodes will know the answer for humans (Figure 6E). Notably, both transitive inference and sensory preconditioning require hippocampus (Dusek and Eichenbaum, 1997; Gilboa et al., 2014; Wikenheiser and Schoenbaum, 2016), entorhinal cortex (Buckmaster et al., 2004) and ventral prefrontal cortex (Jones et al., 2012; Koscik and Tranel, 2012). For example, manipulations to any of these structures will preserve an animal’s preference for A over B or B over C, but abolish the preference for A over C (even though A has been rewarded many times more than C) (Figure 6B).

**Figure 6:**
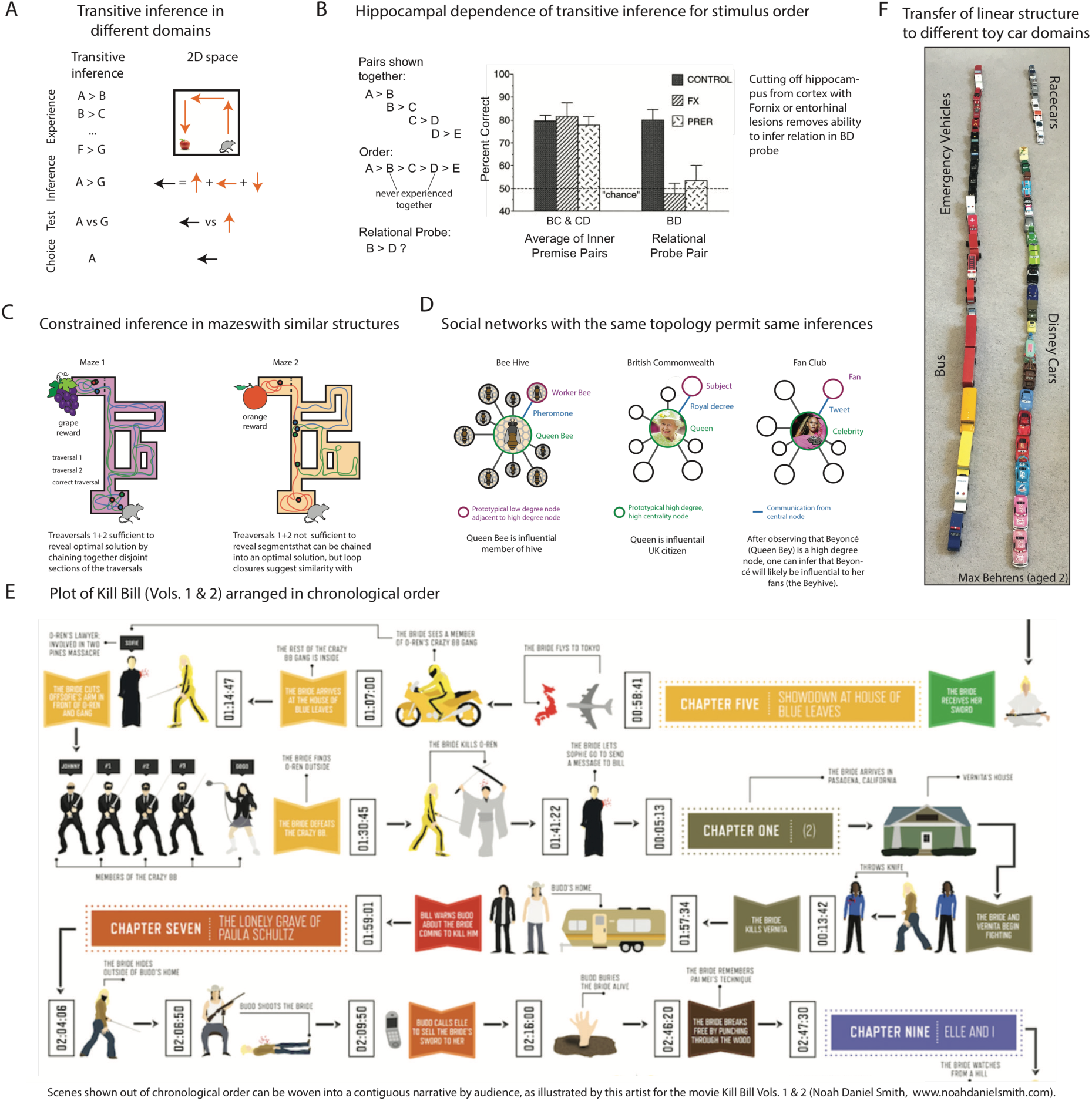
Transitive inference and structural constraints on inference. A) Breakdown of experience, inference, and choice at test time in Spatial navigation and transitive inference both rely on chaining together separately observed sequences of observations. B) (Dusek and Eichenbaum, 1997) show that transitive inference is hippocampally dependent in rats. Rats were trained on pairs of consecutive stimuli, in which the rewarded stimulus on each trial followed the order A>B>C>D>E. If the animal represents the stimulus order rather than simply the results of the experienced pairs, it can correctly infer that B should be preferred over D. Control animals make this inference correctly, while animals with lesions that separate hippocampus from cortex are impaired on these relational probes. C) Illustration of Mazes that permit different types of structured inferences. In Maze 1, neither of the first two trajectories traverse the shortest path solution; however, pieces from the traversals can be joined to compose the shortest path. In similarly structured Maze 2, the observed trajectories are insufficient to compose the shortest path. However, the loop closures suggest topological similarity to the previously observed Maze 1. This similar previously experienced relational structure can be used to constrain inference in the new maze with a representation that factors observation from underlying structure. D) Prior experience with topological features can be used to constrain inference in non-spatial environments as well. For instance, exposure to the abstract notion of a “high-degree node,” which arises in many different types of social networks, can be used to draw conclusions about the role of a high-degree individual in a novel network. For instance, a queen bee, the Queen of England, and Beyoncé (“Queen Bey”) are members of very different social networks, but fill a similar role of influence relative to other individuals in the network. E) Humans can fluidly chain together episodes experienced out of order into a continuous narrative. This is elegantly illustrated by an artist’s depiction of the Plot of Kill Bill Vols 1&2, arranged in chronological order (Noah Daniel Smith, www.noahdanielsmith.com). Although scenes of Kill Bill are seen out of order, such that early scenes depicting some event are qualified by later scenes depicting an earlier event, a viewer can integrate these scenes into a coherent narrative. Other films with this property include Memento and Pulp Fiction (http://www.noahdanielsmith.com/pulpfiction/). (F) “Lines of cars” by Max Behrens aged 2. Organisation within each line suggests that structures can be placed within other structures. This organisation is readily visible in the left-hand line, but will be most apparent in the right-hand line to readers with domain expertise in the Disney Film Series “Cars”.

#### What experiences have in common

Why should the brain learn general structural representations rather than build a new representation for each task? For this to be a useful strategy, there must be regularities in the world that can be profited from. And indeed there are – the world is brimming with repetition and self-similarity at every level of abstraction (Figure 6). Knowledge can be generalised about objects and concrete entities – if you find a fish in a lake, it’s worth checking other lakes for fish; or about transition structures – multiple rooms often lead from the same corridor; or about the relationship between objects and transitions – if you reach a sad point of a film, you are probably about half way in. Repetitions exist in the relationships between objects – if two people are friends on Facebook, they probably follow similar people on Twitter (Figure 6C-D). Crucially, the structures that organise these self-repetitions often themselves repeat across nature (Kemp and Tenenbaum, 2008). Tree-like organisation, for example, can be found in families, in rumour-mills, and even in trees. “Small-world” and “scale-free” properties are found in complex systems across the natural world (Watts and Strogatz, 1998).

Across life, then, a learner faces a *distribution of tasks* (Figure 7A), and this distribution is not random but highly structured. Each new task can be constrained by rich prior information from previous tasks. Harlow’s “learning set” is a clear controlled example. In Harlow’s experiment, what is randomized between episodes is the identity of each object; while what is constant is the relationship between object and reward. Harlow’s interpretation of “Learning to learn” was that past experience drove the acquisition of abstract structure – for example, the fact that “one of the two objects is always rewarded” – and this learned representation made future learning more efficient.

**Figure 7:**
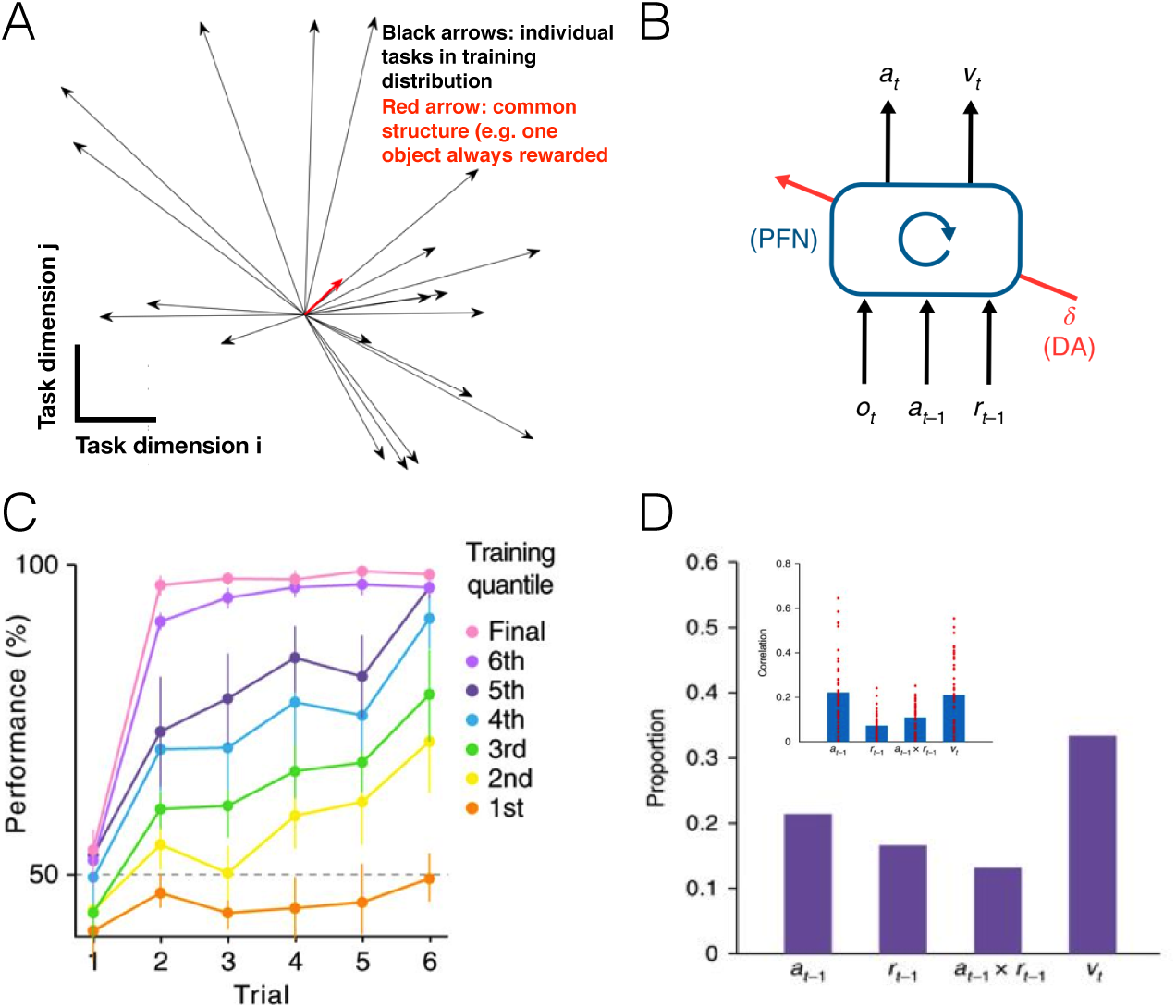
Meta-learning and meta-reinforcement learning. A) A learner faces a distribution of tasks that share common underlying structure. Consider again the Harlow task. In any individual episode the actual sensory-motor-reward contingencies are such that there are many possible algorithms that would work to obtain reward. For example, “always choose the blue object”, or “choose the left object unless the left object is different from last time and the square object is on the right”, etc (black arrows). However, the only thing that is common across all of the different task episodes is “one of the two objects is always rewarded” (red arrow). This common structure is what meta-learning seeks to acquire, to facilitate future learning. With a slow enough learning rate, such that individual task episodes do not dominate the learning, the result is that a solution can be learned that works for all tasks. B) One method for performing meta-learning is meta-reinforcement learning (meta-RL). The meta-RL architecture from (Wang et al., 2018) consists of a prefrontal network (PFN) modelled as a recurrent neural network (RNN) with synaptic weights adjusted through an RL algorithm (driven by dopamine, DA). At each time step, the agent receives the current observation *o* and the past action *a* and reward r, which are fed into a recurrent neural network that outputs an action and state value estimate *v*. This RNN learns to use its activation dynamics as a second free-standing RL algorithm, adapted to the task distribution on which it is trained. C) Through training, meta-RL learns to learn faster on Harlow’s task, similar to the monkeys in Figure 2B. Adapted from (Wang et al., 2018). D) Meta-RL provides a possible explanation for how individual units in PFC become tuned to various task-related variables. Proportion of recorded neurons in macaque PFC with sensitivity to last action, last reward, action-by-reward interaction, and current value (Tsutsui et al., 2016). Inset: for each unit in the trained meta-RL agent, strength of correlation with the same variables. Adapted from (Wang et al., 2018).

Recent approaches in artificial neural networks have demonstrated that, with enough experience, powerful and general structural representations can emerge from simple principles. In the following sections we will highlight some of these principles that seem particularly relevant to the neuronal representations and anatomical constraints that can be found in the frontal cortex and hippocampal formation.

#### Learning Structure from experience

Deep learning techniques can learn powerful representations of tasks that closely resemble biology (Mante et al., 2013; Sussillo et al., 2015; Yamins and DiCarlo, 2016), and there are several approaches for adapting these techniques to learn structural knowledge. These are collectively referred to as meta-learning (Andrychowicz et al., 2016; Finn et al., 2017; Hochreiter et al., 2001), but one with a tantalizing link to brain function is meta-Reinforcement Learning (meta-RL) (Wang et al., 2018) (Box 1).

Meta-RL solves the reinforcement learning problem (maximise expected task reward) with a recurrent neural network (RNN) whose weights are trained through a reward prediction error signal (Figure 7B). The critical insight, however, is that the network does not need to change its weights to react to rewards and errors in solving the current task. Such reactions can instead be encoded in the dynamics of the network – because an RNN can provably implement any algorithm (Siegelmann and Sontag, 1995), it can implement an RL algorithm if it is trained to do so. The network weights are then trained to maximise reward over many different tasks by setting a learning rate that is too slow to accommodate learning within a single task, but appropriate to average experience across many tasks.

The consequence of this, is that the reward prediction error signal drives the weights to encode the structure that is common across episodes, rather than information about any particular sensory inputs. The activation dynamics of the network can then profit from this structure to produce rapid learning in each task. For example, in Harlow’s task, like the human and non-human primates, after many training episodes the network can learn to solve the problem in one trial (Figure 7C). In other experiments, Wang et al also found that meta-RL learned abstract notions about the dynamics of the environment, independent of its current state, and used these to learn more efficiently from new experience.

This observation provides a potential solution to an intriguing conundrum in the neuroscience of reinforcement learning. Dopamine signals a reward prediction error that is assumed to cause learning but, in prefrontal cortex at least, triggers synaptic change over the wrong timescale – tens of seconds at the fastest (Brzosko et al., 2015; Otmakhova and Lisman, 1996; Wang et al., 2018; Yagishita et al., 2014) when behaviour can change within a few seconds or less (i.e., the animal gets a reward and immediately changes its behaviour). Meta-RL proposes that instead of driving learning directly, the role of dopamine is to drive changes in the *learning algorithm;* this learning algorithm is implemented in the recurrent circuits particularly centring on the prefrontal cortex.

Consistent with this, when the meta-RL agent is trained on typical reward learning tasks, individual units in the network acquire tuning properties that resemble individual PFC units recorded in monkeys in the same task. For example, in a foraging task, (Tsutsui et al., 2016) found a variety of units in macaque PFC, with some coding predominantly for value, others for previous action, others for reward, and still others for an action-by-reward interaction. When meta-RL was trained to perform the same task, individual units in the artificial network spontaneously acquired tuning for these variables, with a similar distribution to the monkey neurons (Tsutsui et al., 2016)(Figure 7D).

#### Factorisation and constraints – how should structural knowledge be represented?

It is of course possible to represent relationships between objects in an implicit fashion – encoded in the synaptic weights between object representations. For example, in sensory preconditioning, one can easily imagine scenarios in which cells that form the representation of object A form new synaptic connections to those that encode object B. Indeed, such mechanisms likely do exist in the brain (e.g. in (Grewe et al., 2017)). However, in order for a structural abstraction (such as the linear order in transitive inference and sensory preconditioning, or the 2D-layout of physical- and bird-space) to generalise from one task to another, its representation must be **explicit** – divorced from the sensory properties of the particular task in question, and should be in a form that allows it to impose its constraints on any new sensory environment (Box 1). One way to enforce such a representation mathematically, is to require representations to *factorise*, such that the probability distribution of activity for a task event is the product of two independent distributions defining the sensory and structural contributions to the task. Mathematically this is:

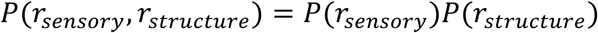

where the probability is, for example, defined over the spiking (**r**) of each neuron in the representation. Factorisation facilitates learning because it dramatically reduces the dimensionality of the representation to be learnt, and allows extreme forms of generalisation outside of the training data. For example, if you want to predict how your daughter will react to a blue cup you can learn the blue distribution from all blue things (not just cups) and the cup distribution from all colours (not just blue) (Figure 8A). Using these two independent distributions, you can predict conjunctions you have never experienced (Figure 8B). Similarly, if the representation of a line is factorised from the representation of the elements of that line it can be learnt across many different tasks and generalised to new ones.

**Figure 8:**
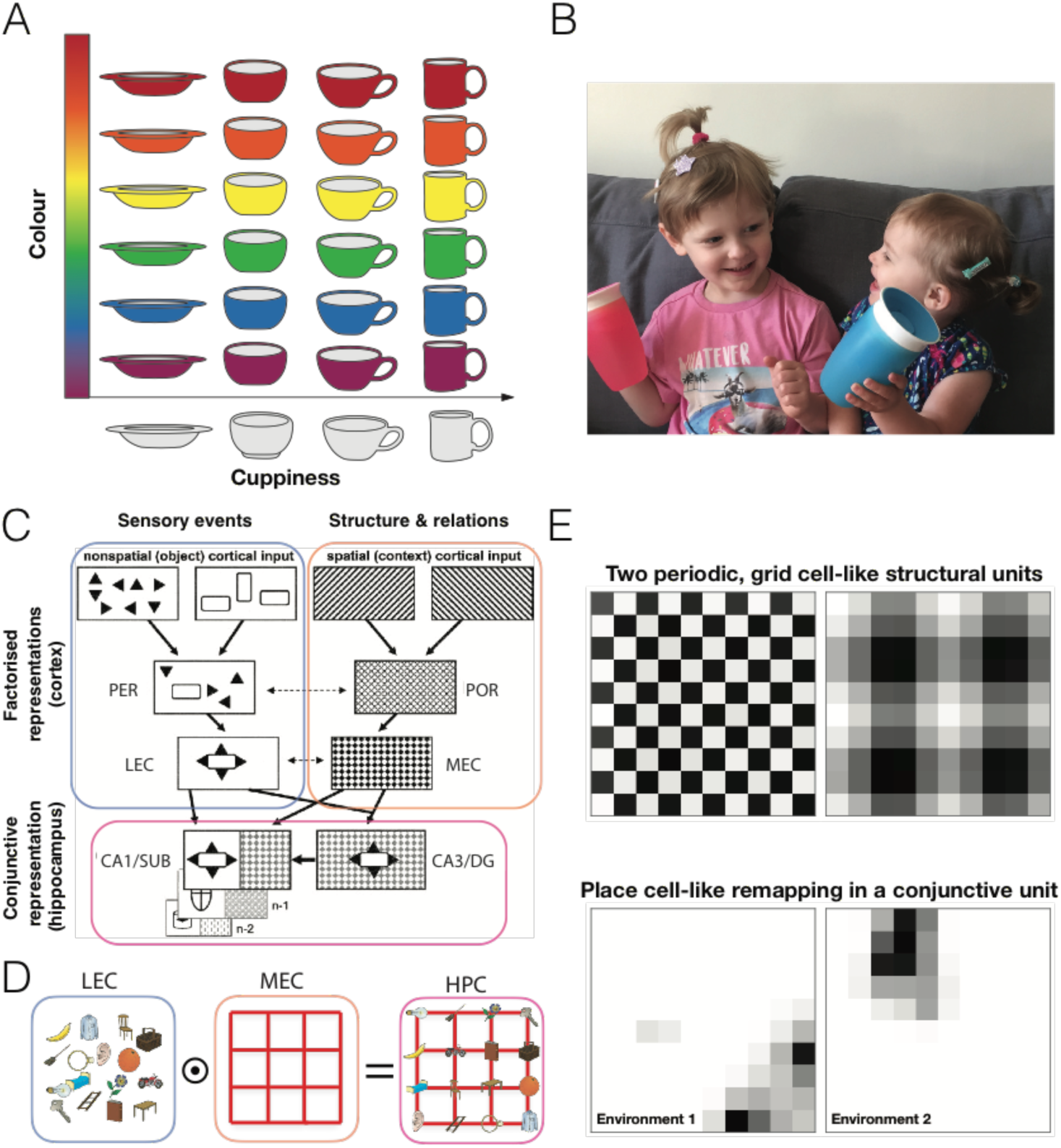
Factorised and conjunctive codes. A) Entities in the world can be factorised in to independent distributions defining the dimensions of these entities. For example, different objects may be factorised according to how ‘cuppy’ they are as well as their colour. B) Factorisation allows generalisation to as yet unseen conjunctions. For example, it is easy to imagine young children with unusual colour preferences, such as Max and Lana (Min) Behrens. C) The functional representations in the hippocampal-entorhinal system are suggestive of factorisation and conjunction. In the hippocampal inputs, medial regions code structure devoid of sensory information, lateral regions code sensory information devoid of structure. These are combined in a conjunctive code in hippocampus. Figure adapted from (Manns and Eichenbaum, 2006). HPC = hippocampus, MEC = medial entorhinal cortex, LEC = lateral entorhinal cortex, PER = perirhinal cortex, POR = postrhinal/parahippocampal cortex. D) Denotes this separation in a simple task environment containing a graph with different objects at the vertices. E) An artificial neural network that explicitly encourages factorised and conjunctive codes learns units with properties similar to grid and place cells. Units representing the structure of the environment, in this case a 2D topology, have periodic firing fields similar to grid cells (a square rather than triangular lattice is learned due to a four way connected space). Units coding for the conjunction between the learned structure and sensory events exhibit remapping, similar to place cells. Adapted from (Whittington et al., 2018).

These considerations are intriguing when considering the cellular representations that can be found in the hippocampal formation and its inputs (Manns and Eichenbaum, 2006) (Figure 8C). In the brain regions that precede hippocampal activity by one or a small number of synapses, representations are separated (factorised) into structural (spatial/contextual) representations in medial regions and sensory (object) representations in lateral regions. By contrast hippocampus proper contains conjunctive representations of object in structure. Cells are only active for a particular object in a particular location but not for the object or location alone (Komorowski et al., 2009; Wood et al., 1999). Whilst the conjunctive hippocampal representation is sufficient to fully represent the current episode (importantly for memory), cortical regions that summarise the statistics of these episodes (McClelland et al., 1995), do so in a factorised, data efficient manner that can be conjoined in hippocampus to represent episodes that have never been experienced.

By modelling such a factorised system in a neural network, it is possible to examine the properties of the structural representations (Whittington et al., 2018). Just like the rodents we discussed earlier, when tasked with predicting the next sensory event in a 2D random-walk, the network can profit from knowledge of the 2D spatial structure; for example, it can know (/infer) such things as if I go up, left, down and right I will be back in the same place. This allows for correct predictions of sensory events when re-visiting states, even when approaching from an entirely new direction – knowing the structure means knowing where you are in the space. To do this the network must learn a factorised representation of the structure. This structural representation can then be combined with sensory events in a conjunctive code to form different memories at different places in different rooms. The learnt structural representations include periodic cells (similar to grid cells) (Figure 8D), but also cells that resemble boundary cells and boundary vector cells. Since sensory events may occur in different locations in different rooms, the conjunction between a given sensory event and the structural representation may be in a different location across rooms. Therefore the conjunctive units naturally exhibit remapping (Figure 8D), analogous to the firing of place cells in different environments. It is possible, then, that at least some of the cell-types that make up the rich spatial representation in the hippocampal formation can be accounted for by structural considerations that generalise to arbitrary non-spatial problems.

#### Structural bases and the hippocampal zoo

We have argued that repeating structural constraints on tasks should be embedded explicitly in the neural code, and that entorhinal cells provide examples of such a representation. For simple tasks (such as reversal learning or object discrimination), it is possible for the brain to represent the exact transitions from one state to the next, but this strategy will break as soon as anything changes in this structure. For example, the appearance of a shiny object dramatically changes the transition statistics of all tasks if you are a magpie. More prosaically although boundaries always have the same effect on transitions, they might do so in different parts of the state space in different environments. One solution to this problem is to think of the entorhinal cells as a basis set (Box 1) for describing the current transition structure, so that different combinations of cellular activity can represent the different structural constraints of different environments (similar to how cells in primary visual cortex represent a basis for describing the pixel distributions in natural images (Olshausen and Field, 1996)). These bases will capture common features across tasks and whilst the most reliable of these features will be most strongly represented (such as the translational and scale invariance of grid cells), there will also be more minor representations of features that have less prominent structural influence. Indeed, recent evidence suggests that activity in many entorhinal cells that are hard to classify into easily interpretable cell types, are nevertheless linearly predictive of task-relevant behavioural variables (Hardcastle et al., 2017). When starting a new task, the recurrent neural network in the Entorhinal cortex may represent an initial guess at the task structure, from which the hippocampus can form conjunctive codes and memories. With task experience, a more appropriate weighting of the basis functions can be inferred, and thus task structure more correctly approximated (Barry et al., 2012). This interpretation is consistent with the strong attractor dynamics apparent in the grid cell network (Burak and Fiete, 2009) as grid cells embed the relationships that are the most prevalent amongst spatial tasks. Given known hippocampal involvement in primate behaviours with markedly different statistical structure from space (such as social tasks), it is possible that other statistical features are also similarly deeply embedded.

#### What should be built by evolution, and what should be learnt from your environment?

Learning about structure and assuming constraints are highly complementary. The more abstract (further from observations) a structure is, the harder it is to learn (Raghu et al., 2016), but such abstractions can be easily hard-coded by evolution. For example, the factorisation of relationships from objects in the cortical code immediately bestows the abstract principle that “Different pairs of objects might have relationships in common”. To obtain such an abstract bias with meta-learning alone can be extremely difficult because it’s hard to make such a diverse task distribution that this principle is the only thing the tasks have in common. Indeed, modern machine learning techniques are exploring how to hard code these abstract biases into artificial neural networks. For example, the current state of the art in Starcraft II (a multiplayer real-time strategy game set in a distant part of the Milky Way) was achieved by adding a relation network component that employs exactly this principle (Zambaldi et al., 2018). By contrast, meta-learning is better suited for discovering complex biases that are difficult to program directly, and which may be unexpected properties of task families of interest.

These two different strengths play well together. Starting with abstract architectural biases can make other, more specific or complex biases easier to meta-learn (Zambaldi et al., 2018), and can encourage more generalisable structural representations to emerge. Indeed, the representations learned in the factorised network described above (Whittington et al., 2018) are basis functions (as are those found in (Banino et al., 2018)), which generalise to environments of different sizes. Pure meta-learning without any inductive biases, however, may require a broader task distribution to learn such generalisable representations. Thus, evolution should provide architectural biases that facilitate generalisable structure learning.

It is still unclear how much of the bias observed in the brain – like a belief in relational structure – is completely hard-coded versus learned through early life experience. It is of course possible that particular structural constraints that apply broadly across natural tasks (such as 2D maps, ordinal lines, hierarchies) have been favoured on evolutionary timescales, and are therefore hardwired into cortical connectivity. Such an argument might explain the precise anatomical arrangement of grid cell modules along the dorsal-ventral axis of entorhinal cortex (Brun et al., 2008).

### Implications for the cognitive map

In the remainder of this review, we would like to take a more speculative position and consider what implications the structural abstractions we have discussed above might have on the “systematic organisation of knowledge” that Tolman envisaged. This organisation clearly encompasses much more than the representation of structural abstractions, but these structures constrain how concrete objects and actions will be combined. These constraints not only allow objects to be configured into meaningful current or future events, but also provide a powerful means to generalise learning from sparse observations.

#### Inferential planning

In both machine learning (Kocsis and Szepesvári, 2006; Silver et al., 2016; Sutton and Barto, 1998) and neuroscience research (Daw et al., 2005, 2011), it is often assumed that planning your future involves searching through a tree of possible states and discovering the best one. This process is so costly as to be impossible in most reasonable circumstances, so an alternative is to estimate a cached value of states that are your immediate neighbours or neighbours to some depth (Huys et al., 2015; Keramati et al., 2016). Armed with structural knowledge, however, it is possible that plans can be built into representations in an analogous fashion to the representation of objects on transitive-inference lines. The planning process now becomes an inference of what should go where on the line. This inference is further constrained by domain-specific structural knowledge of relationships between items. When planning a lab, a PI does not search through every possible arrangement, but instead knows that they should hire the theorist before the experimentalist to avoid wasted experiments, and the experimentalist before the data scientist to avoid twiddled thumbs. By constraining the representations of the different objects (postdocs!) by this relational knowledge, the number of possible futures are limited. If similar structural knowledge (A relies on B) has been used in previous plans such as building a house (foundations, walls, windows) then all that is necessary to build the new plan is for the object’s representation to contain information about which other objects this structural knowledge applies to. The new plan can then be generalised (inferred) from the old. This idea is an extension of ideas in psychology, where object representations are assumed to encompass possible actions that the object ‘affords’ (Gibson, 1966). The representation of the word “apple” for example, might include activity that represents the facial movements required to eat it, and the hand position required to grip it. In this way, infinite possible actions are reduced to only a few likely ones conditioned on the objects currently available.

Placed together with structural and relational knowledge, however, such representations would provide dramatic constraints on possible long-term futures and powerful generalisations to novel scenarios. Whilst there is little experimental evidence along these lines to date, it is notable that what evidence there is centres on the hippocampus and ventromedial prefrontal cortex. These regions are active during both reconstructive memory and constructive imagination (Buckner and Carroll, 2007) and without these regions people can’t construct imagined futures (Hassabis et al., 2007). Indeed, when subjects imagine the taste of a new food that they have never experienced but is constructed from known ingredients (such as Tea-Jelly), both regions show evidence that cellular ensembles for the ingredients are active simultaneously (Barron et al., 2016).

The examples above give a sense of the power of combining different structural representations (here lines and reliances) and of the same structural representations being generalised across domains. The logic is similar in spirit to arguments made about the compositional nature of human visual understanding, where known elements can be composed into new objects that can be immediately understood with no prior experience (Lake et al., 2015, 2017). This analogy, however, highlights the importance of a feature of structural coding emphasised in earlier sections. In order for structures to play an analogous role in compositional planning as objects do in compositional vision, they cannot be encoded solely in the synaptic weights between object representations. They must be represented explicitly, as are the objects that they act upon. Indeed this blurring of the line between of objects and structures is a powerful feature of human cognition, allowing us to reason about structures and relationships. A marriage, for example, is a concrete event, or a structure for organising our social knowledge, or a profound set of constraints on future behaviour.

#### Structural inferences for generalised learning

It is possible, then, for futures to be inferred (or generalised) rather than planned. By corollary, sparse observations can cause profound learning when constrained by structural knowledge. When reports of Austrian nobles crossing the Pyrenees reached Louis XIV of France, he was able to use the same structural relation (A relies on B – here wedding relies on guests) to infer a proposed alliance between the Spanish and Holy Roman Empires. His subsequent plan to detain the bride-to-be at Versailles (inferred, presumably from the same relationship applied to brides rather than guests) led to the War of the Spanish succession (at least in the BBC’s interpretation (BBC, 2018)). By filtering experiences through a scaffold of relational knowledge, precise inferences can be drawn from little data (Lake et al., 2015).

If structures and relations are represented explicitly, however, there is the potential for inference at dramatic levels of abstraction. The social experience of watching a parent cajoling a child to new bravery, for example, can be replicated even when people or animals are replaced with abstract shapes (Heider and Simmel, 1944). The dynamic relationship between the two triangles on the screen is sufficient for us to infer the roles of parent and child and the motives and emotions of each. As with the non-spatial grid cells described earlier, a structure (albeit a more complex and dynamic one) that is evolved or learnt to describe the behaviour in one setting, can be generalised to a completely different domain. Whilst it might seem initially mysterious why there is an evolutionary benefit to infer social dynamics between triangles, or 2 dimensions of birds, it is clear that the ability to infer structural analogies between disparate situations has profound consequences for learning. Getting saved by a bicycle helmet might, for example, make you more likely to wear a safety jacket next time you are on a boat, or to take out home insurance next time you are at your computer. In modern artificial intelligence research, there is a substantial effort to discover learning rules that ensure “continual learning” – that is learning rules that enable new tasks to be learnt by networks without destroying old ones (Kirkpatrick et al., 2017; Zenke et al., 2017). In our view, structural abstractions and inferences are a key element to this endeavour.

It takes a particularly dramatic form of selective attention to be a cognitive neuroscientist. When a subject walks into the laboratory, reads a complex set of instructions and effortlessly translates them into a complex sequence of future events and actions to be performed inside a 13-tonne sarcophagus that they happily enter because a stranger has told them it is safe, it takes an unusual degree of restraint to choose to study neural activity when the same subject receives payments that differ by 15 pence. When animals in the wild are capable of building sophisticated networks of burrows, for example, or allegiances, it takes a similar degree of focus to choose to study how they navigate an open 1m square, or whether they prefer one stimulus to another after several months of training. This selective attention has, however, been profitable because it has allowed experiments to be performed in a theoretical framework where they can build on one-another in a formal sense. We envisage that the nascent emergence of formalisms to describe more complex, flexible behaviours will, similarly, provide a framework for profitable experiments in this broader behavioural sphere, and further encourage the exciting re-emergence of collaborations between protagonists of artificial and biological intelligence.

##### Box 1

###### States of the world

A state is a possible configuration of the world. Whilst the true state of the world is complex and high dimensional, only a very small part of this state is relevant to the animal performing a task. The animal therefore has a **state definition problem**. If they can give different names to states in which these **relevant** dimensions are different but the same names to states which only differ in **irrelevant** dimensions, they will dramatically reduce the learning problem, whichever learning algorithm they employ. For example, a waiter learning efficient strategies for opening wine bottles should have separate states for corks vs screw-tops but not for red vs white contents. This way they will learn fastest because, with two states rather than one, they will be able to learn two different strategies and, with two states rather than four, they will have twice as much experience on each strategy.

###### Latent states

Problems arise when these relevant dimensions are not **observable**. Whilst waiters can often see the tops of their wine bottles, there is little immediate sensory data to tell a driver whether they can or cannot use the bus route. One solution to this problem is for the driver to build different **latent states** that allow different routes to be planned during commuting and non-commuting times.

###### Models of the world

A **model** of a world is an internal representation the world’s structure that can be used to predict future states of the world. A good model encompasses a parsimonious **state definition** but also an understanding of how **each state transitions to the next**. This is usually expressed as **p**(**s’| s,a**), the probability that each state will transition to each other state if you choose action **a**. If you have such a model you can clearly plan your future, but you can also learn more efficiently as the model places strong constraints on possible explanations of sensory data. If the cake is burnt, it is unlikely to be due to the extra spoonful of sugar.

**Learning set** describes accrued knowledge from prior tasks that allows for stereotyped learning on new tasks. If an animal arrives at a new task equipped from prior tasks with a model of the task, or even with a parsimonious state representation, they will learn the new task much faster. In Harlow’s example, this learning set came from prior experience of the exact same task, but it is clear that knowledge can be generalised across different tasks that share features (Figure 2F). Faced with a screw-driver, it is likely advantageous to have learnt to open a bottle.

**Meta-learning** is the machine learning term for learning set, whereby features that are common amongst many tasks are exploited to speed up learning on a particular task instance. There are a number of approaches to meta-learning. One example is **meta**-**reinforcement learning** (meta-RL) in which meta-learning is driven by reward prediction errors. More general forms of meta-learning might also be driven by sensory observations. In machine learning, meta-learning can be used to tune synaptic weights, but also to tune neural architectures resulting in interesting parallels with learning over the lifespan and adaptation over evolutionary timescales respectively.

###### Factorised representations

For a learning set, separating the structural representation from the representation of sensory particularities of the task facilitates reusing the structural representation in related tasks. Factorisation is one way to achieve this separation. Here the probability distribution of the task is a product of two independent distributions describing the model/structural and sensory contributions:

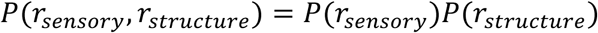

Having learned both component distributions, flexible generalisation can be made to entirely new task conjunctions, for example a different arrangement of sensory events (Figure 8A-B). Such representations are reminiscent of the separation of structural from sensory information in the cortices surrounding hippocampus (Figure 8C), and are consistent with the reuse of structural information across different rooms in remapping experiments in medial entorhinal cortex (but not hippocampus) (Figure 3A).

###### Basis representations

If learning set can act across different tasks, then the representation of the model cannot be a hardcoded state diagram. It must be flexible such that new tasks can be constructed from combinations of elements of old tasks. These are termed bases and should represent common features of tasks in a manner that allows flexible recombination. This broad definition encompasses bases for how features should be combined into a **state definition** (different bottle tops should imply different states) but also bases for common patterns of **transition structure** in the world (1D, 2D, tree etc), and for how **state features impact on this transition structure** (animals tend to approach interesting objects, and are unlikely to walk through walls). These last two types of bases appear to have commonalities with entorhinal cell populations.

###### Eigenvector basis

One important basis representation is the eigenvector basis, as it is the linear basis that explains the most variance in the data per basis function (or cell!) and is therefore a particularly efficient way to represent the structure of a task. An interesting recent observation is that the eigenvectors of the transition function of 2D space have strong similarities to grid cells (Figure 5).

###### Structural inference

When encountering a new task, animals should infer what structural knowledge they should use to guide their decisions. They can do this either by inferring structure by observed sensory features (in a restaurant, it is likely that the eating will follow the sitting), or inferring structure from observed transitions (what is next in the sequence 2,4,6,8,10,… ?), or a combination. In reinforcement learning, the inferred structure can be used to define the possible state spaces and constrain the possibilities of relations between different states while learning the state space of a new task. In this paper, we suggest that having an explicit representation of structure (in the form of structural bases) can help in solving such inference problems and therefore in learning new state spaces.

## Acknowledgments

Thanks to Thomas Akam and Philipp Schwartenbeck for very helpful comments on the paper. We acknowledge funding from Wellcome Trust Senior Research Fellowship (WT104765MA) together with a James S. McDonnell Foundation Award (JSMF220020372) to T.E.J.B.

## Footnotes

1 There are important differences between different regions of the ventral frontal cortex and we are aware that, by ignoring them here, we invite scepticism and irritation from our colleagues in equal measures. Given the existing complexity of the argument we are putting forward, we ask to be excused our anatomical imprecision and refer readers to the following papers for interesting discussions of these differences (Haber and Behrens, 2014; Rudebeck and Murray, 2014; Rushworth et al., 2011). It is perhaps worth noting here that, in our experience, one difference that predicts whether activity (or lesion effects) will be most prominent medially or laterally within the ventral frontal cortex pertains to whether it reflects a computation that is used to guide choices (medially), or a computation that is used to learn from the outcomes of those choices (laterally).

